# Competing Fluid Forces Control Endothelial Sprouting in a 3-D Microfluidic Vessel Bifurcation Model

**DOI:** 10.1101/626531

**Authors:** Ehsan Akbari, Griffin B. Spychalski, Kaushik K. Rangharajan, Shaurya Prakash, Jonathan W. Song

## Abstract

Sprouting angiogenesis, the infiltration and extension of endothelial cells from pre-existing blood vessels, helps orchestrate vascular growth and remodeling. It is now agreed that fluid forces, such as laminar shear stress due to unidirectional flow in straight vessel segments, are important regulators of angiogenesis. However, regulation of angiogenesis by the different flow dynamics that arise due to vessel branching, such as impinging flow stagnation at the base of a bifurcating vessel, are not well understood. Here we used a recently developed 3-D microfluidic model to investigate the role of the flow conditions that occur due to vessel bifurcations on endothelial sprouting. We observed that bifurcating fluid flow located at the vessel bifurcation point suppresses the formation of angiogenic sprouts. Similarly, laminar shear stress at a magnitude of ∼3 dyn/cm^2^ applied in the branched vessels downstream of the bifurcation point, inhibited the formation of angiogenic sprouts. In contrast, co-application of ∼1 µm/s average transvascular flow across the endothelial monolayer with bifurcating fluid flow and laminar shear stress induced the formation of angiogenic sprouts. These results suggest that transvascular flow imparts a competing effect against bifurcating fluid flow and laminar shear stress in regulating endothelial sprouting. To our knowledge, these findings are the first report on the stabilizing role of bifurcating fluid flow on endothelial sprouting. These results also demonstrate the importance of local flow dynamics due to branched vessel geometry in determining the location of sprouting angiogenesis.

## Introduction

Blood vessels comprise a hierarchical network that transports oxygen and nutrients throughout the body (Pries and Secomb, 2014). Expansion of this network occurs by angiogenesis, where endothelial cells (ECs) that line the inner surface of all blood vessels, sprout and branch to support tissue nourishment and growth (Carmeliet and Jain, 2011; Potente et al., 2011). Angiogenesis is necessary to help repair injured tissue (Greaves et al., 2013), while uncontrolled angiogenesis is a prominent characteristic of rapidly growing solid tumors (Kerbel, 2008). Therefore, the growth and remodeling of blood vessels by angiogenesis is critical throughout physiology (Virmani et al., 2005; Cao, 2007), and increasing our fundamental understanding of angiogenesis is necessary to improve therapeutic strategies used for regenerative medicine, cardiovascular medicine, and cancer therapy.

Research on angiogenesis has traditionally focused on identifying biochemical signaling factors that regulate EC function (Shibuya, 2013). Yet in physiology, blood vessels are continuously exposed to fluid mechanical forces due to blood flow, including shear stress tangential to the endothelium, and transverse flow across the vessel wall. Emerging research has highlighted the importance of the forces created by blood flow in potently influencing the angiogenic process. These studies in the *in vitro* setting have been buoyed largely by the advancements in microfabrication techniques that enable the development of perfusable models that integrate 3-D tissue scaffolds for investigating angiogenesis in response to controlled fluid forces (Wong et al., 2012; Akbari et al., 2017). For instance, application of 3 dyn/cm^2^ laminar shear stress (LSS) was shown to suppress endothelial sprouting induced by vascular endothelial growth factor (VEGF) (Song and Munn, 2011), while LSS values greater than 10 dyn/cm^2^ have been shown to induce angiogenic sprouting (Galie et al., 2014). Furthermore, previous studies have shown the pro-angiogenic role of transvascular flow (TVF) that is driven by a transmural pressure difference between the vasculature and the adjacent interstitium (Song and Munn, 2011; Vickerman and Kamm, 2012; Galie et al., 2014).

Endothelial sprouting has predominantly been examined in *in vitro* microfabricated devices with straight blood vessel models. However, the vasculature *in vivo* consists of a hierarchy of branching structures that generates bifurcating fluid flow (BFF) at the base of vessel bifurcations. BFF can be characterized by local stagnation pressure imparted normal to the endothelium and near-zero average shear stress, distinguishing BFF from previously studied hemodynamic factors (i.e., LSS and TVF). Recently, we reported an *in vitro* microfluidic model of a branching vessel that enables measurement of vascular permeability under application of BFF, LSS, and TVF (Akbari et al., 2018) while also presenting an interface between fluid flow, ECs, and 3-D extracellular matrix (ECM). The findings from this study demonstrated the vascular permeability outcomes in response to the local flow dynamics at specific locations along the bifurcating vessel structure. However, the role of BFF in co-regulating endothelial sprouting has not been reported previously. Since elevated vascular permeability typically coincides with increased angiogenesis (Nagy et al., 2012), the purpose of this paper was to investigate endothelial sprouting due to the flow dynamics produced by branching vessel geometry.

Here we report that application of BFF with ∼38 dyn/cm^2^ stagnation pressure at the vessel bifurcation point (BP) and ∼3 dyn/cm^2^ LSS in each branched vessel (BV) inhibits the formation of angiogenic sprouts. Furthermore, co-application of TVF with BFF and LSS induces the formation of angiogenic sprouts, thereby suggesting the presence of competing effects between TVF with BFF and LSS in regulating endothelial sprouting in a vessel bifurcation model. This work presents the first quantitative report on the stabilizing role of BFF for inhibiting endothelial sprouting. Moreover, these results demonstrate that sprout location can be determined by the local flow dynamics due to vessel bifurcations. Consequently, the findings reported here advance the current understanding of the fluid mechanical regulators of angiogenesis.

## Materials and Methods

### Microfluidic model of vessel bifurcation

A microfluidic platform was developed as a 3-dimensional (3D) *in vitro* analogue of a bifurcating vessel as previously described (Akbari et al., 2018) (Fig. 1A, B). The microchannels were fabricated by soft lithography of polydimethylsiloxane (PDMS). Upon seeding, the microchannels were fully lined with mouse aortic endothelial cells (MAECs) to form an *in vitro* vessel analogue (Fig. 1D). An important feature of the microfluidic model is a 3-D ECM compartment comprised of a mixture of collagen gel (3 mg/ml rat tail-type I) and fibronectin (10 μg/ml) that was positioned between the two parallel BV regions and intersects the base of the branching vessel or the BP (Fig.1D). Located at the BP and two BV’s are 100 µm wide openings (referred to as apertures) between two PDMS posts that help contain the pre-polymerized collagen gel solution in the 3-D ECM compartment (Huang et al., 2009). These apertures enabled direct contact of the MAEC monolayer with the supporting collagen ECM where sprouts can infiltrate (Fig. 1E).

**Figure 1 –.**
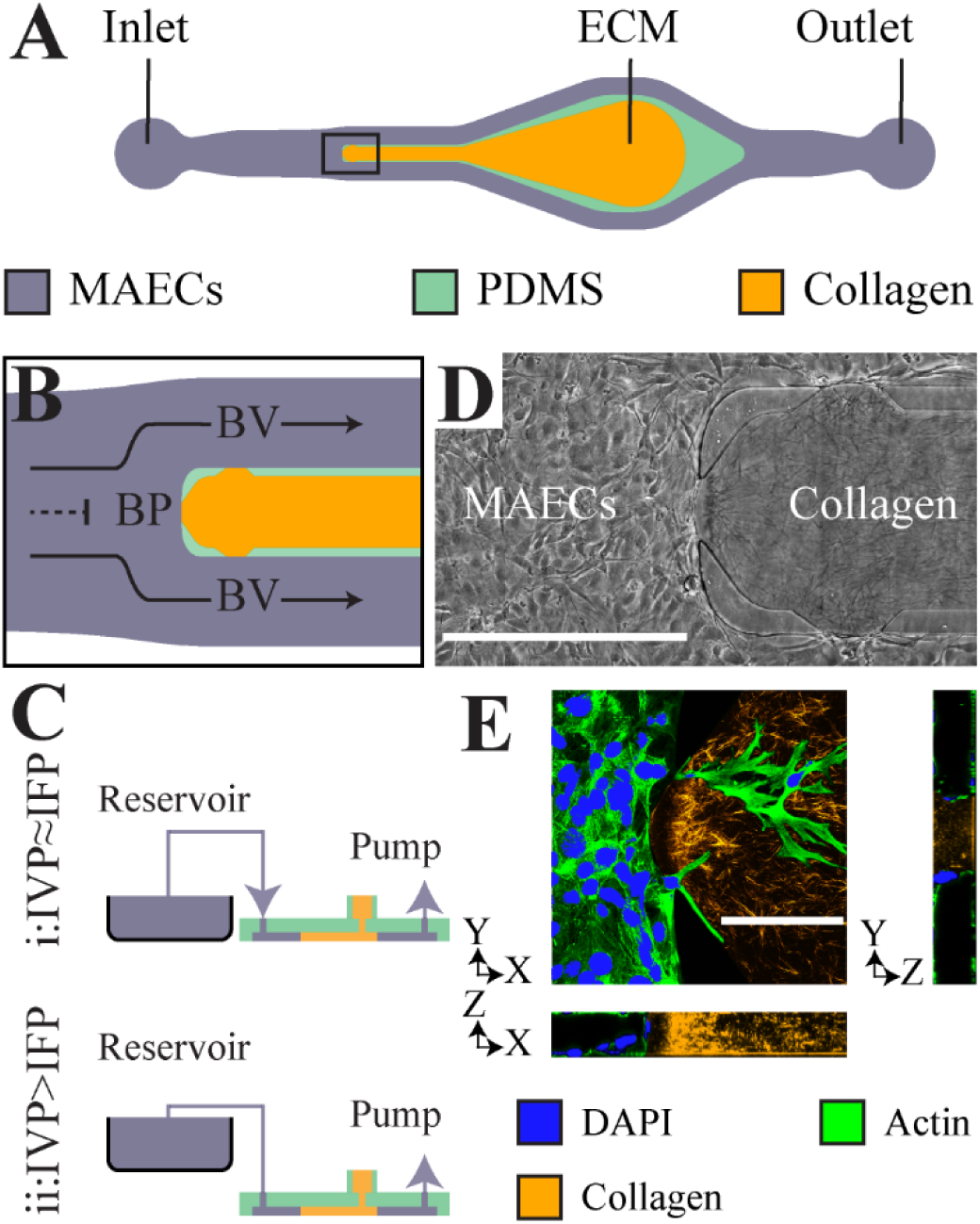
Schematic and characterization of the *in vitro* microfluidic device reported previously (Akbari et al., 2018) used here for examination of sprouting angiogenesis. (A) Top-view for the microfluidic device. (B) Schematic showing a close-up for the local fluid flow at the BP aperture and the flow along apertures in the BV. (C) The schematic of the approach implemented to control the difference between IVP and the IFP through controlled elevation of the reservoir. (D) A representative phase-contrast image of the BP fully lined with MAECs adjacent to the polymerized ECM hydrogel. Scale bar, 500 μm. (E) A representative confocal image of the microfluidic device fully seeded with MAECs. Furthermore, the 3D structure of the polymerized collagen matrix (orange) was resolved using total reflectance confocal imaging. Scale bar, 100 μm.

### Cell culture and perfusion of the microfluidic model

MAECs were generously provided by the laboratory of Dr. Mike Ostrowski and cultured as previously described (Srinivasan et al., 2009). Briefly, MAECs were grown in DMEM-F12 cell culture media supplemented with 20% heat-inactivated fetal bovine serum, 1% penicillin-streptomycin, 10 U/mL heparin and 30 μg/mL endothelial cell growth supplement. The microchannels were pre-treated with fibronectin (10 μg/ml) for 2 hours at 37°C followed by treatment with cell culture growth media overnight at 37°C. Subsequently, MAECs at 6-10 passage number were seeded at 20,000 cells/μl concentration, allowed to adhere overnight at 37°C, and grown to confluence for 24 hours. Controlled perfusion in the microfluidic model was applied using a programmable syringe pump as previously described (Akbari et al., 2018). A full-scale computational model of the microfluidic platform was developed using COMSOL Multiphysics (version 4.4) to determine: a) the average LSS experienced by the MAECs that are adhered on the ECM at the BP and in each BV, b) the level TVF resulting in interstitial flow based on the measured hydraulic permeability of the ECM and the endothelial cell monolayer (Akbari et al., 2018).

### Immunofluorescence

Following treatment with each experimental condition, the microdevices were flushed 3 times with phosphate buffered saline (PBS) and fixed using 3% paraformaldehyde for 30 min at room temperature. Next, the devices were flushed 3 times with PBS and incubated with the blocking buffer (5% goat serum with 0.1% Triton X-100 in PBS) for 1 hour at room temperature. The devices were then flushed 3 times with PBS and incubated with phalloidin (solution made by diluting the stock by 1:20 in PBS with 10% blocking buffer) for 30 min at room temperature to stain for actin. Next, the devices were flushed 3 times with PBS and incubated with DAPI (solution made by diluting the stock by 1:1000 in double distilled water) for 5 min at room temperature. Finally, the devices were flushed 3 times with PBS prior to epifluorescence imaging (TS-100, Nikon).

### Quantification of increase in sprouting area

Sprouting area was quantified using NIH ImageJ. A user-defined region of interest was defined for each aperture (i.e. BP or BV) and the total area covered by sprouting MAECs was quantified. The total sprouting area at 72 hours after seeding was subtracted by the sprouting area at 24 hours after seeding to report the total increase in sprouting area over 48 hours.

### Quantification of cellular elongation and alignment

The morphological responses of MAEC sprouting were quantified using an elongation index and angle of orientation, which are commonly used parameters that quantify the extent that cells elongate and align in the direction of flow, respectively (Song et al., 2005). The elongation index for each sprouting MAEC was defined by the ratio between the major and minor axis of an ellipse fit to the cell area using MATLAB. The angle of orientation between each sprouting MAEC and interstitial flow was defined by the absolute angle between the major axis of the ellipse fit to the cell area and the interstitial flow vector at the cell area centroid that was identified by DAPI nuclear stain. An angle of orientation of 0 degrees is a cell aligned perfectly with the direction of interstitial flow and 90 degrees is a cell aligned perpendicular to the direction of interstitial flow.

### Statistical analysis

Each experimental test condition was conducted at least in triplicate to report statistical analysis. The values reported for the sprouting area represent the average ± standard error of the mean. Two-sided student t-test was used to report statistical significance between each two pair of test conditions. The following symbols were used to report statistical significance: * indicates *p*-value < 0.05, *** indicates *p*-value < 0.001.

## Results

### Bifurcating fluid flow (BFF) and laminar shear stress (LSS) attenuate endothelial sprouting

The branching vessel geometry of our 3-D microfluidic model enables evaluation of angiogenic sprouting in response to BFF and LSS within the same fluidic circuit. In addition, our microfluidic model enables control of TVF levels independent of the perfusion rate, thereby enabling us to decouple the effects of TVF on endothelial sprouting from the effects of BFF and LSS. This capability is achieved by controlling the pressure difference via a static pressure head column between the IVP and interstitial fluid pressure (IFP) domains (Fig. 1C). Consequently, TVF can be either induced or restricted based on whether there is a difference in IVP and IFP (Akbari et al., 2018). When IVP is equal to IFP (IVP=IFP), TVF is ∼0 (Fig. 2A). Thus, under these experimental conditions, sprouting angiogenesis responses were due to BFF and LSS and in the absence of TVF. When IVP is greater than IFP (IVP > IFP), a 1.5 cm H_2_O hydrostatic pressure difference is produced that results in a TVF of ∼1 µm/s oriented from the MAEC-lined intravascular region, across the endothelium, and into the interstitial ECM compartment (Fig. 2A). Under these conditions, sprouting angiogenesis was evaluated in response to co-application of BFF and LSS alongside TVF.

**Figure 2 –.**
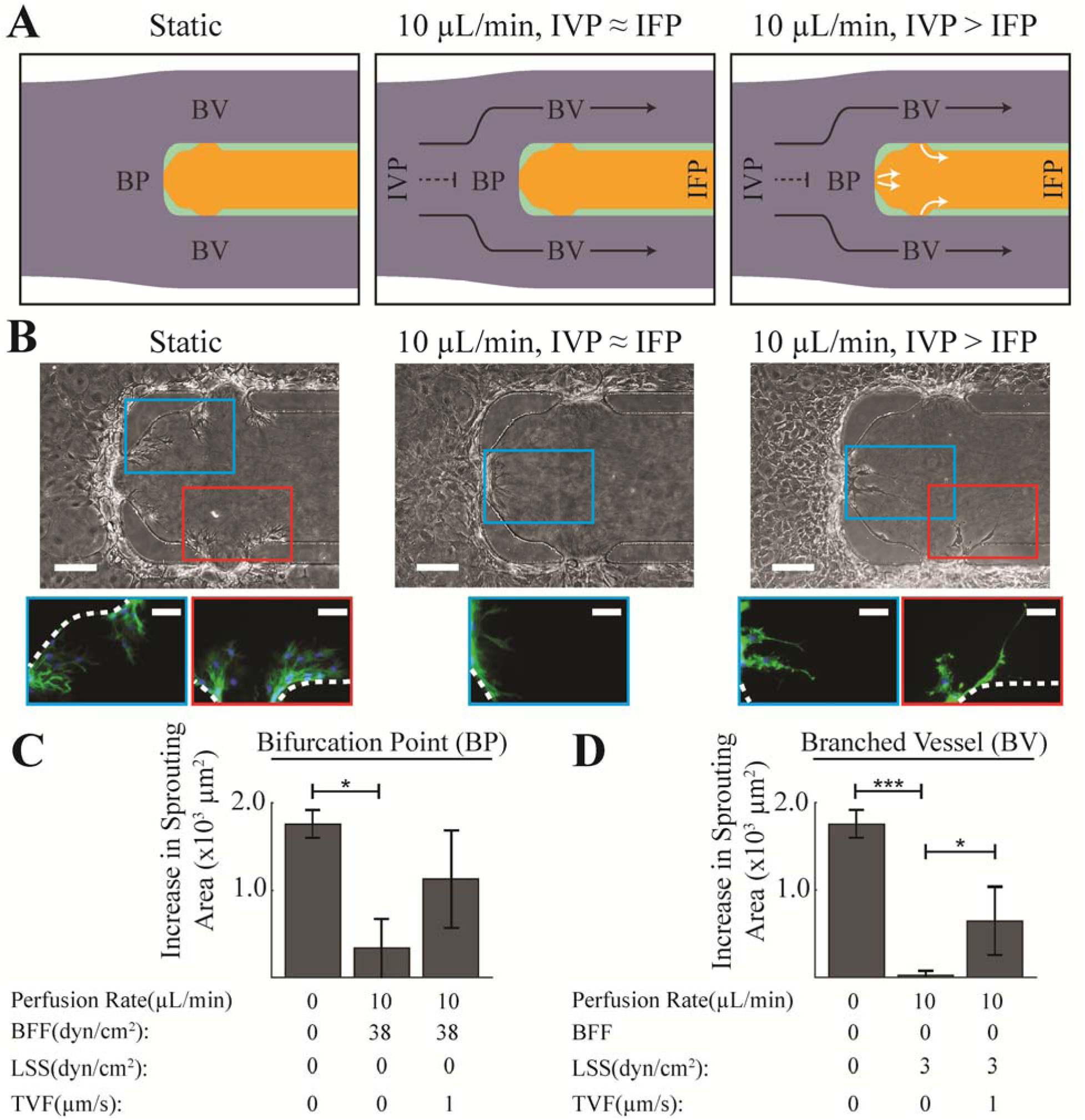
Application of (BFF and LSS attenuated endothelial sprouting. (A) The top-view schematics of the microdevice depicting the fluid mechanical factors under: (i) static, (ii) perfusion with equilibrated IVP and IFP resulting in BFF (dashed black line) at the BP, LSS (solid black line) in each BV and negligible TVF, and (iii) perfusion with elevated IVP resulting in BFF at the BP, LSS in each BV and 1µm/s TVF (white solid lines). (B) Representative phase contrast and epifluorescence images of sprouting MAECs in response to perfusion with ∼10 μL/min after 48 hours under equilibrated and elevated IVP compared to static control condition. Dashed white lines mark the location of the PDMS posts. (C, D) Quantitative representation of the level of increase in sprouting area in response to treatment with each experimental test condition: at the BP, and in BV. *: *p_value* < 0.05. ***: *p_value* < 0.001.

The microfluidic devices were perfused with complete MAEC growth medium at a volumetric flow rate of 10 µl/min to generate ∼3 dyn/cm^2^ LSS in BV as previously described (Akbari et al., 2018). This LSS level is within the physiological range of shear stress in the microcirculation *in vivo* (Aird, 2005). In addition, previously reported computational estimations showed that 10 µl/min perfusion flow rate in this microfluidic model generates ∼38 dyn/cm^2^ stagnation pressure at the BP imparting a normal force due to stagnation flow at the BP along with near-zero shear stress (Akbari et al., 2018). Application of ∼38 dyn/cm^2^ BFF for 48 hours, in the absence of TVF, significantly attenuated endothelial sprouting area by ∼80% compared to the static control condition where MAECs undergo spontaneous sprouting (Fig. 2B and 2C). Similarly, treatment with ∼3 dyn/cm^2^ LSS for 48 hours, in the absence of TVF, completely inhibited sprouting area (i.e. reduced by ∼98% compared to the static control condition) (Fig. 2B and 2D). The observed suppression of MAEC sprouts by ∼3 dyn/cm^2^ LSS is in line with previously reported results *in vitro* with human umbilical vein endothelial cells (HUVECs) in response to approximately the same LSS levels (Song and Munn, 2011). However, to our knowledge, our findings are the first report that endothelial sprouting is attenuated by BFF.

### TVF competes with BFF and LSS to induce formation of angiogenic sprouts

Once establishing the effects of BFF and LSS on sprouting, next we investigated the effects of systematically introducing TVF (∼1 µm/s) at the fixed flow rate of 10 µl/min that produces ∼38 dyn/cm^2^ BFF and 3 dyn/cm^2^ LSS in our bifurcation model. Co-application of TVF with BFF at the BP resulted in a ∼3-fold increase in MAEC sprouting area compared to BFF alone, although this response was not statistically significant (Fig. 2B and 2C). Similarly, co-application of TVF with LSS at the BV regions resulted in a ∼28-fold increase in MAEC sprouting area compared to LSS alone (Fig. 2B and 2D). The observed competing effects between TVF versus BFF and LSS on sprouting is in accordance with previous reports on the pro-angiogenic effect of TVF (Song and Munn, 2011; Galie et al., 2014). Furthermore, the level of endothelial sprouting induced by co-application of ∼1µm/s TVF alongside BFF and LSS was lower compared to the level of sprouting under static control condition. These findings suggest that the observed stabilizing effects of 38 dyn/cm^2^ BFF and 3 dyn/cm^2^ LSS counteracts the vessel sprouting responses triggered by ∼1 µm/s TVF.

### Interstitial flow streamlines coordinate elongation of the angiogenic sprouts

Once establishing the effects of BFF, LSS, and TVF in triggering or suppressing endothelial sprouting, next we investigated whether the interstitial flow through the 3-D ECM compartment due to IVP>IFP (Fig. 2A) supported the alignment and elongation of MAEC sprouts. It is known that endothelial cells cultured in 2-D align and elongate with the direction of LSS within 24 hours (Levesque and Nerem, 1985). In addition, it was previously shown that when exposed to interstitial flow, breast cancer cells aligned parallel to flow streamlines (Polacheck et al., 2011). The MAEC sprouts that formed in response to co-application of TVF alongside BFF and LSS at the BP and each BV, respectively, were more elongated (elongation index of 5.9±0.9) compared to static control condition (elongation index of 2.4±0.2). These results suggest that interstitial flow through the ECM helps promote the extension of sprouting MAECs.

Next, we examined whether the local interstitial streamlines can help guide the direction of MAEC sprouting. Interstitial flow streamlines that were previously determined by computational modeling (Akbari et al., 2018) were superimposed onto the optical microscopy images of MAEC sprouting into the 3-D ECM (Fig. 3). At the BP, MAEC sprouts that formed by co-application of TVF and BFF aligned with the direction of the local interstitial flow streamlines, as demonstrated by the angle of orientation measurement of 6.8±3.3 degrees (Fig. 3A). In contrast, MAEC sprouts at the BV and formed under co-application of LSS and TVF do not prominently align with the local interstitial flow streamlines (angle of orientation = 48.6±5.4 degrees) (Fig. 3A). These outcomes were expected because at the BP aperture, the direction of TVF and interstitial flow in the supporting ECM are approximately parallel. In contrast, at the BV, the direction of interstitial flow changes prominently within ∼50 μm away from the aperture interfaces (Fig. 3A). Therefore, the MAEC sprout that extend from the BV aperture interfaces are required to reorient in order to align with the direction of interstitial flow. Finally, we confirmed that in the absence of MAECs, collagen matrix fiber orientation was not altered by perfusion and interstitial flow over 48 hours (Fig. 3B). Thus, the observed alignment between angiogenic sprouts and the corresponding local interstitial flow streamlines is not mediated by flow-induced reorientation of the collagen fibers.

**Figure 3 –.**
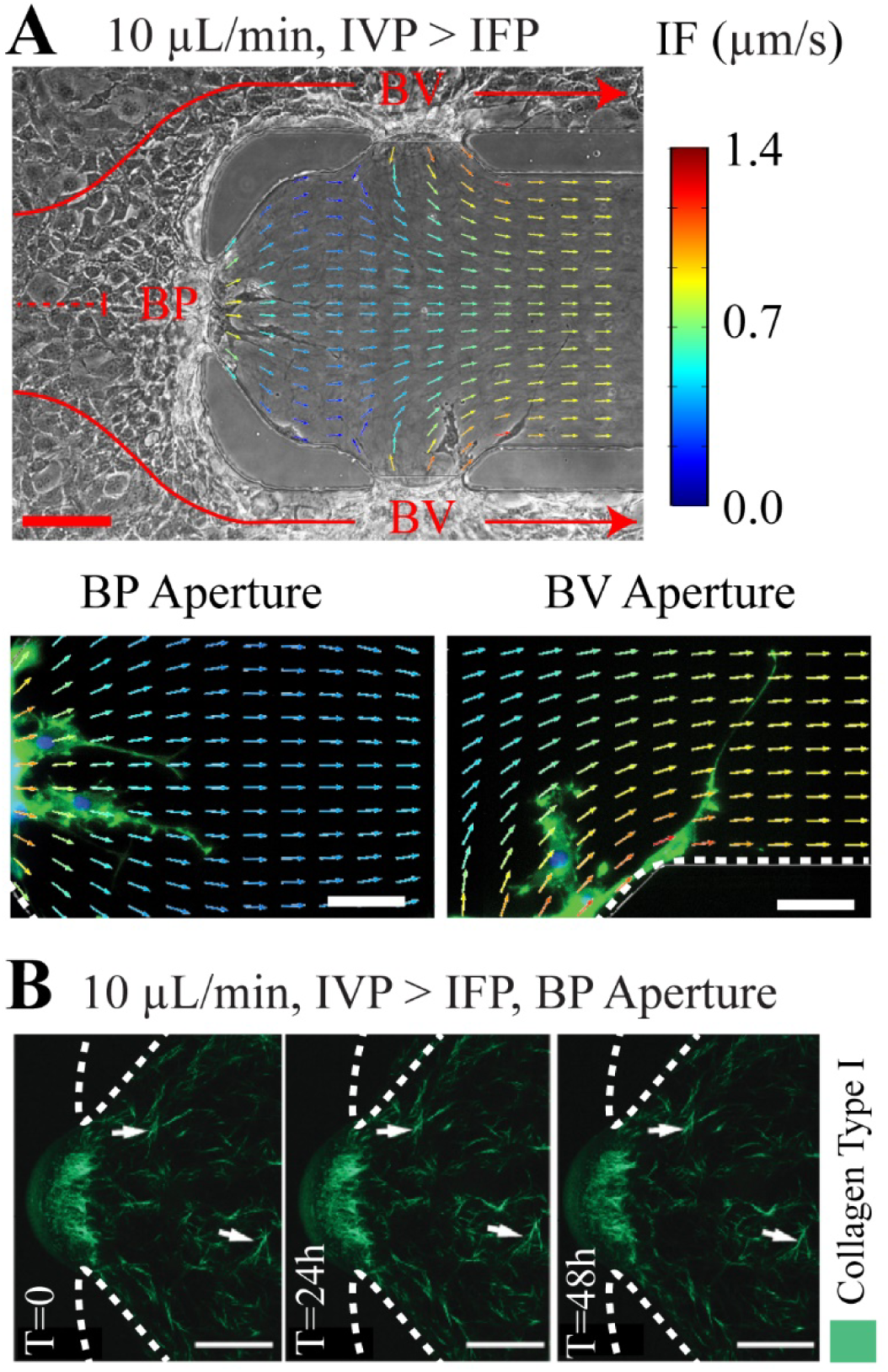
The angiogenic sprouts formed under elevated IVP were more elongated and showed alignment with the interstitial flow streamlines. (A) The computational interstitial flow streamline map superimposed on representative phase contrast and epifluorescence images of the angiogenic sprouts formed under elevated IVP condition. (B) Confocal reflectance microscopy images of collagen fibers under application of interstitial flow in the absence of MAECs. Arrowheads in the panels indicate landmark fibers to track among these images. Interstitial flow did not elicit a direct change in collagen fiber orientation. Red scale bar, 100 µm. White scale bars, 50 µm. Dashed white lines mark the location of the PDMS posts.

## Discussion

Obtaining a detailed and mechanistic understanding of the main physical regulators of angiogenic vascular sprouting and remodeling is significant for advancing therapeutic strategies for modulating pathological angiogenesis. These studies can be facilitated by biomimetic platforms that reconstitute tissue level function *in vitro* while also enabling controlled application of pressure and shear stress to cultured endothelial cells. In this work, the effect of stagnation pressure at vessel bifurcations and LSS on sprouting angiogenesis in presence and absence of TVF was evaluated. The studies were conducted in an *in vitro* microfluidic device that presents BFF in the presence of a collagen ECM. The results reported here show, for the first time, that endothelial sprouting was inhibited at the bifurcation point at 38 dyn/cm^2^ stagnation pressure with nearly no shear stress. Application of 3 dyn/cm^2^ LSS in the BV also imparted a stabilizing effect on the endothelium by suppressing the formation of angiogenic sprouts compared to static control conditions. While the LSS results agree with the previously reported role for tangential fluid forces in inhibiting VEGF induced endothelial sprouting (Song and Munn, 2011), and decreasing endothelial permeability (Buchanan et al., 2014; Akbari et al., 2018) the inhibition of sprouting due to a stagnation pressure is a new finding. It is worth noting that past *in vivo* observations on increased endothelial sprouting from vessels with low to no blood flow such as damaged or occluded vessels (Yamada et al., 1999) or blind-ending sprouts (Anderson et al., 2008) also corroborate the reported stabilizing effect of 3 dyn/cm^2^ LSS.

At the BP, the BFF represents negligible LSS but significant stagnation pressure arising from an impinging fluid flow. Yet, BFF did not show uncontrolled angiogenic sprouting. A previous *in vivo* study in the context of embryonic development showed that angiogenic sprouts form from points with local minimum shear stress except when the minimum shear occurs at the convergence of two blood vessels (Ghaffari et al., 2015). Interestingly, the fluid mechanical conditions at vessel convergences also show similar flow characteristics of possible flow stagnation based on the included angle of the incoming streams and the relative fluid shear of the converging fluid streams (Prakash et al., 2009). Furthermore, it was recently demonstrated that 38 dyn/cm^2^ BFF at the BP for 6 hours induced a significant decrease in endothelial permeability, thus causing stabilization of the endothelium (Akbari et al., 2018). In other words, as BFF stabilizes the endothelium, it is logical to expect a reduction the sprouting angiogenesis, which is the outcome we observed in this study.

Interestingly, simultaneous co-application of transvascular flow (TVF) alongside 38 dyn/cm^2^ BFF and 3 dyn/cm^2^ LSS induced the formation of angiogenic sprouts at BP and in each BV, respectively. Therefore, while BFF and LSS stabilize the endothelium, the TVF competes with the effects of BFF and LSS to induce formation of angiogenic sprouts. The sprouts formed under co-application of TVF alongside BFF and LSS were more elongated compared to static control condition. Furthermore, the angiogenic sprouts formed at the BP under elevated IVP (i.e., presence of BFF and TVF) was visually observed to be along the direction of interstitial flow streamlines at the BP, but not at the BV (i.e., presence of LSS and TVF) after 48 hours. The visual alignment observations also find supporting evidence when taken together with the previous results on the pro-angiogenic effect of fluid flow across the endothelial monolayer (Vickerman and Kamm, 2012; Galie et al., 2014).

It is noteworthy that we observed endothelial sprouting with the direction of TVF and interstitial flow. Numerous studies using other microfluidic models have reported that angiogenic sprouts preferentially form against the direction of interstitial flow (Song and Munn, 2011; Song et al., 2012; Vickerman and Kamm, 2012; Kim et al., 2016). However, these previous studies were done with HUVECs, which are venous microvascular ECs, in contrast to our study using MAECs, an arterial endothelial cell type. In addition, a previous study demonstrated that LSS inhibited sprouting in venous or capillary endothelial cell types (i.e. HUVECs and HMECs) but not arterial endothelial cells (BAECs) (Tressel et al., 2007). Therefore, one must consider the cell source when interpreting the results of sprouting phenotypes due to flow dynamics.

Previous studies have attempted to discover the primary endothelial mechanosensory pathway that transduces the mechanical stimuli from LSS to the downstream endothelial biological response (Davies, 1995; Hahn and Schwartz, 2009). Since BFF is characterized by near-zero average shear stress at the stagnation point, it is plausible if the mechanotransduction pathway through which BFF leads to endothelial response is distinct from the studied pathways for LSS. Therefore, the detailed signaling pathways pertaining to BFF mechanotransduction requires further investigations. Moreover, while our results were in the context of the blood vessel morphogenesis, it has been observed that intraluminal lymphatic valves preferentially form at the bifurcation points of collecting lymphatic vessels (Udan and Dickinson, 2012). Our microfluidic model can be readily adapted to study the morphogenetic events coordinated by the local flow dynamics of branched vessels with applications for both the blood and lymphatic vasculature.

In summary, the results reported here introduce BFF as a potent regulator of endothelial sprouting while showing that both bifurcating flows and tangential shear flows are important determinants of angiogenic sprouting in an *in vitro* microfluidic model of vessel bifurcations.

## Acknowledgements

Funding support was provided by The American Heart Association (15SDG25480000), NHLBI (R01HL141941), and Center for Emergent Materials, an NSF-MRSEC grant DMR-1420451. Images presented in this report were generated using the instruments and services at the Campus Microscopy and Imaging Facility, The Ohio State University. This facility is supported in part by grant P30 CA016058 from the NCI. Partial personnel support through the US Army Research Office through grant number W911NF-16-0278 is also acknowledged. E.A. and K.K.R acknowledge OSU Presidential Fellowships. G.B.S. acknowledges funding from the Undergraduate Summer Research Program of The American Heart Association Great Rivers Affiliate, a Barry M. Goldwater Scholarship, and an Ohio State University (OSU) Comprehensive Cancer Center Pelotonia Fellowship.

## References

Aird, W.C. (2005). Spatial and temporal dynamics of the endothelium. J Thromb Haemost 3(7), 1392-1406. doi:10.1111/j.1538-7836.2005.01328.x.

Akbari, E., Spychalski, G.B., Rangharajan, K.K., Prakash, S., and Song, J.W. (2018). Flow dynamics control endothelial permeability in a microfluidic vessel bifurcation model. Lab Chip 18(7), 1084-1093. doi:10.1039/c8lc00130h.

Akbari, E., Spychalski, G.B., and Song, J.W. (2017). Microfluidic approaches to the study of angiogenesis and the microcirculation. Microcirculation 24(5). doi:10.1111/micc.12363.

Anderson, C.R., Hastings, N.E., Blackman, B.R., and Price, R.J. (2008). Capillary sprout endothelial cells exhibit a CD36 low phenotype: regulation by shear stress and vascular endothelial growth factor-induced mechanism for attenuating anti-proliferative thrombospondin-1 signaling. Am J Pathol 173(4), 1220-1228. doi:10.2353/ajpath.2008.071194.

Buchanan, C.F., Verbridge, S.S., Vlachos, P.P., and Rylander, M.N. (2014). Flow shear stress regulates endothelial barrier function and expression of angiogenic factors in a 3D microfluidic tumor vascular model. Cell adhesion & migration 8(5), 517–524.

Cao, Y. (2007). Angiogenesis modulates adipogenesis and obesity. J Clin Invest 117(9), 2362-2368. doi:10.1172/JCI32239.

Carmeliet, P., and Jain, R.K. (2011). Molecular mechanisms and clinical applications of angiogenesis. Nature 473(7347), 298-307. doi:10.1038/nature10144.

Davies, P.F. (1995). Flow-mediated endothelial mechanotransduction. Physiol Rev 75(3), 519-560. doi:10.1152/physrev.1995.75.3.519.

Galie, P.A., Nguyen, D.H., Choi, C.K., Cohen, D.M., Janmey, P.A., and Chen, C.S. (2014). Fluid shear stress threshold regulates angiogenic sprouting. Proc Natl Acad Sci U S A 111(22), 7968-7973. doi:10.1073/pnas.1310842111.

Ghaffari, S., Leask, R.L., and Jones, E.A. (2015). Flow dynamics control the location of sprouting and direct elongation during developmental angiogenesis. Development 142(23), 4151-4157. doi:10.1242/dev.128058.

Greaves, N.S., Ashcroft, K.J., Baguneid, M., and Bayat, A. (2013). Current understanding of molecular and cellular mechanisms in fibroplasia and angiogenesis during acute wound healing. J Dermatol Sci 72(3), 206-217. doi:10.1016/j.jdermsci.2013.07.008.

Hahn, C., and Schwartz, M.A. (2009). Mechanotransduction in vascular physiology and atherogenesis. Nat Rev Mol Cell Biol 10(1), 53-62. doi:10.1038/nrm2596.

Huang, C.P., Lu, J., Seon, H., Lee, A.P., Flanagan, L.A., Kim, H.Y., et al. (2009). Engineering microscale cellular niches for three-dimensional multicellular co-cultures. Lab Chip 9(12), 1740-1748. doi:10.1039/b818401a.

Kerbel, R.S. (2008). Tumor angiogenesis. N Engl J Med 358(19), 2039-2049. doi:10.1056/NEJMra0706596.

Kim, S., Chung, M., Ahn, J., Lee, S., and Jeon, N.L. (2016). Interstitial flow regulates the angiogenic response and phenotype of endothelial cells in a 3D culture model. Lab Chip 16(21), 4189-4199. doi:10.1039/c6lc00910g.

Levesque, M.J., and Nerem, R.M. (1985). The elongation and orientation of cultured endothelial cells in response to shear stress. J Biomech Eng 107(4), 341-347. doi:10.1115/1.3138567.

Nagy, J.A., Dvorak, A.M., and Dvorak, H.F. (2012). Vascular hyperpermeability, angiogenesis, and stroma generation. Cold Spring Harb Perspect Med 2(2), a006544. doi:10.1101/cshperspect.a006544.

Polacheck, W.J., Charest, J.L., and Kamm, R.D. (2011). Interstitial flow influences direction of tumor cell migration through competing mechanisms. Proc Natl Acad Sci U S A 108(27), 11115-11120. doi:10.1073/pnas.1103581108.

Potente, M., Gerhardt, H., and Carmeliet, P. (2011). Basic and therapeutic aspects of angiogenesis. Cell 146(6), 873-887. doi:10.1016/j.cell.2011.08.039.

Prakash, S., Akberov, R., Agonafer, D., Armijo, A.D., and Shannon, M.A. (2009). Influence of Boundary Conditions on Sub-Millimeter Combustion. Energy & Fuels 23(7), 3549-3557. doi:10.1021/ef900040q.

Pries, A.R., and Secomb, T.W. (2014). Making microvascular networks work: angiogenesis, remodeling, and pruning. Physiology (Bethesda) 29(6), 446-455. doi:10.1152/physiol.00012.2014.

Shibuya, M. (2013). Vascular endothelial growth factor and its receptor system: physiological functions in angiogenesis and pathological roles in various diseases. J Biochem 153(1), 13-19. doi:10.1093/jb/mvs136.

Song, J.W., Daubriac, J., Tse, J.M., Bazou, D., and Munn, L.L. (2012). RhoA mediates flow-induced endothelial sprouting in a 3-D tissue analogue of angiogenesis. Lab Chip 12(23), 5000-5006. doi:10.1039/c2lc40389g.

Song, J.W., Gu, W., Futai, N., Warner, K.A., Nor, J.E., and Takayama, S. (2005). Computer-controlled microcirculatory support system for endothelial cell culture and shearing. Anal Chem 77(13), 3993-3999. doi:10.1021/ac050131o.

Song, J.W., and Munn, L.L. (2011). Fluid forces control endothelial sprouting. Proc Natl Acad Sci U S A 108(37), 15342-15347. doi:10.1073/pnas.1105316108.

Srinivasan, R., Zabuawala, T., Huang, H., Zhang, J., Gulati, P., Fernandez, S., et al. (2009). Erk1 and Erk2 regulate endothelial cell proliferation and migration during mouse embryonic angiogenesis. PLoS One 4(12), e8283. doi:10.1371/journal.pone.0008283.

Tressel, S.L., Huang, R.P., Tomsen, N., and Jo, H. (2007). Laminar shear inhibits tubule formation and migration of endothelial cells by an angiopoietin-2 dependent mechanism. Arterioscler Thromb Vasc Biol 27(10), 2150-2156. doi:10.1161/ATVBAHA.107.150920.

Udan, R.S., and Dickinson, M.E. (2012). The ebb and flow of lymphatic valve formation. Dev Cell 22(2), 242-243. doi:10.1016/j.devcel.2012.01.022.

Vickerman, V., and Kamm, R.D. (2012). Mechanism of a flow-gated angiogenesis switch: early signaling events at cell–matrix and cell–cell junctions. Integrative Biology 4(8), 863–874.

Virmani, R., Kolodgie, F.D., Burke, A.P., Finn, A.V., Gold, H.K., Tulenko, T.N., et al. (2005). Atherosclerotic plaque progression and vulnerability to rupture: angiogenesis as a source of intraplaque hemorrhage. Arterioscler Thromb Vasc Biol 25(10), 2054-2061. doi:10.1161/01.ATV.0000178991.71605.18.

Wong, K.H., Chan, J.M., Kamm, R.D., and Tien, J. (2012). Microfluidic models of vascular functions. Annu Rev Biomed Eng 14, 205-230. doi:10.1146/annurev-bioeng-071811-150052.

Yamada, H., Yamada, E., Hackett, S.F., Ozaki, H., Okamoto, N., and Campochiaro, P.A. (1999). Hyperoxia causes decreased expression of vascular endothelial growth factor and endothelial cell apoptosis in adult retina. J Cell Physiol 179(2), 149-156. doi:10.1002/(SICI)1097-4652(199905)179:2<149::AID-JCP5>3.0.CO;2-2.

